# scValue: value-based subsampling of large-scale single-cell transcriptomic data for machine and deep learning tasks

**DOI:** 10.1101/2025.01.10.632338

**Authors:** Li Huang, Weikang Gong, Dongsheng Chen

## Abstract

Large single-cell RNA-sequencing (scRNA-seq) datasets offer unprecedented biological insights but pose major computational challenges for visualisation and analysis. Existing subsampling methods can improve efficiency yet may not guarantee downstream machine and deep learning (ML/DL) performance. Here, we propose scValue, a conceptually distinct approach that ranks individual cells by “data value” based on out-of-bag estimates from a random forest. scValue prioritises higher-value cells and allocates more representation to cell types displaying greater value variability, preserving essential biological signals in subsamples. We benchmarked scValue in automatic cell-type annotation tasks on four large datasets (human peripheral blood mononuclear cells, mouse brain cells, human cross-tissue atlas, and mouse aging cell atlas), paired with distinct ML/DL models (scANVI, scPoli, CellTypist, and ACTINN). Our method consistently outperformed existing subsampling methods, closely matching full-data performance in all annotation tasks. Furthermore, in two additional case studies of label transfer learning (via CellTypist) and cross-study label harmonisation (via CellHint), scValue better preserved T-cell annotations across human gut-colon datasets and more accurately reproduced T-cell subtype relationships in a human spleen dataset. Finally, using 16 public datasets ranging from tens of thousands to millions of cells, we compared subsampling quality of scValue and its counterparts on computational time, Gini coefficient, and Hausdorff distance. The method demonstrated fast execution, balanced cell-type representation, and near-random subsampling distributional characteristics. Overall, scValue provides an efficient and accurate solution for subsampling large scRNA-seq data for ML/DL tasks. It is implemented as an open-source Python package installable via pip, with source code available at https://github.com/LHBCB/scvalue.

## Introduction

Large-scale single-cell transcriptomic atlases, such as those from the Human Cell Atlas [1], CELLxGENE [2], SCAR [3], and SCAN [4] provide valuable resources for smaller-scale studies [5]. Visualising and analysing these extensive datasets can pose computational and memory challenges. To address this, researchers can create representative subsets, often referred to as “sketches”, which allow for the extraction of meaningful insights in a more efficient and cost-effective manner [6].

Several sketching methods have been developed so far. As implemented in the Seurat [7] and Scanpy [8] pipelines, uniform subsampling is the simplest method, but may not fully capture the transcriptional diversity of a dataset. To remedy this shortcoming, GeoSketch [6] partitions the gene expression space into equal-sized, non-overlapping boxes and selects representative cells from each box. Meanwhile, Sphetcher [9] covers cells by small-radius spheres that can better preserve the transcriptomic landscape. Both methods enable more evenly distributed sampling across diverse cell types. Hopper [10] provides theoretical guarantees for maintaining the dataset’s structure, while its partition-based variant, TreeHopper, accelerates the sketching process without considerably compromising quality.

GeoSketch, Sphetcher, and (Tree)Hopper all utilises a minimax distance design [11] that minimises the Hausdorff distance, i.e., the maximum distance between any point in the original dataset and the nearest point in the sketch. In contrast, scSampler [12] adopts a maximin distance design [11], maximising an inverse distance measure to ensure that the sampled cells are as distinct as possible. Finally, Kernel Herding (KH) takes a different approach by constructing a “stand-in” sketch [13] that prioritises maintaining the original cell type distribution, rather than optimising the distance metrics.

Machine and deep learning (ML/DL) techniques are becoming increasingly vital in single-cell analysis [14]. However, relatively limited empirical research has focused on evaluating how sketching methods perform in downstream learning tasks. Moreover, assessing the ability of different methods to capture rare cell types, a feature quantified by the Gini coefficient, is an important consideration [12]. In this study, we present scValue, a subsampling method designed to sketch single-cell RNA sequencing (scRNA-seq) datasets based on each cell’s assigned value. Cells deemed to have higher value (i.e., those more informative for cell-type identification) are preferentially included in the sketch, whereas lower-value cells have a reduced likelihood of being selected. Experimental results indicate that scValue outperforms other sketching approaches in ML/DL tasks, yielding more balanced cell-type proportions and exhibiting superior scalability as dataset sizes expand from tens of thousands to millions of cells.

## Materials and Methods

### Large multi-species cross-tissue scRNA-seq datasets

As summarised in Table 1, this study concerns 16 large-scale scRNA-seq datasets containing between 31 thousand and four million cells, spanning four to 197 cell types and including samples from more than 10 tissues across four species. Each dataset was utilised in one or more of the following experiments: demonstration (Demo), ML/DL-based cell type annotation (CTA), label transfer (LT) learning, cell-type harmonization (H), and/or sketch metric comparison (SMC) in terms of computation time, Gini coefficient, and Hausdorff distance.

**Table 1.**
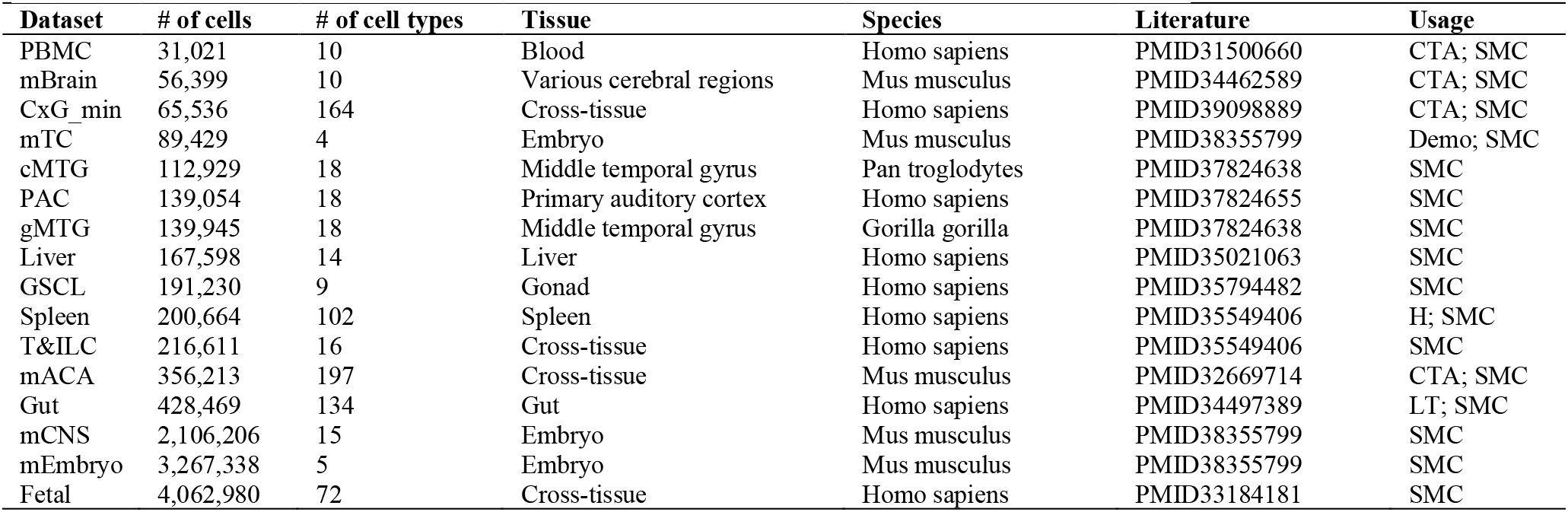
Summary of the 16 scRNA-seq datasets used in this study. Each dataset was utilised in one or more of the following experiments: demonstration (Demo), ML/DL-based cell type annotation (CTA), label transfer (LT) learning, cell-type harmonisation (H), and/or sketch metric comparison (SMC) in terms of computation time, Gini coefficient, and Hausdorff distance.

| Dataset | # of cells | # of cell types | Tissue | Species | Literature | Usage |
| --- | --- | --- | --- | --- | --- | --- |
| PBMC | 31,021 | 10 | Blood | Homo sapiens | PMID31500660 | CTA; SMC |
| mBrain | 56,399 | 10 | Various cerebral regions | Mus musculus | PMID34462589 | CTA; SMC |
| CxG_min | 65,536 | 164 | Cross-tissue | Homo sapiens | PMID39098889 | CTA; SMC |
| mTC | 89,429 | 4 | Embryo | Mus musculus | PMID38355799 | Demo; SMC |
| cMTG | 112,929 | 18 | Middle temporal gyrus | Pan troglodytes | PMID37824638 | SMC |
| PAC | 139,054 | 18 | Primary auditory cortex | Homo sapiens | PMID37824655 | SMC |
| gMTG | 139,945 | 18 | Middle temporal gyrus | Gorilla gorilla | PMID37824638 | SMC |
| Liver | 167,598 | 14 | Liver | Homo sapiens | PMID35021063 | SMC |
| GSCL | 191,230 | 9 | Gonad | Homo sapiens | PMID35794482 | SMC |
| Spleen | 200,664 | 102 | Spleen | Homo sapiens | PMID35549406 | H; SMC |
| T&ILC | 216,611 | 16 | Cross-tissue | Homo sapiens | PMID35549406 | SMC |
| mACA | 356,213 | 197 | Cross-tissue | Mus musculus | PMID32669714 | CTA; SMC |
| Gut | 428,469 | 134 | Gut | Homo sapiens | PMID34497389 | LT; SMC |
| mCNS | 2,106,206 | 15 | Embryo | Mus musculus | PMID38355799 | SMC |
| mEmbryo | 3,267,338 | 5 | Embryo | Mus musculus | PMID38355799 | SMC |
| Fetal | 4,062,980 | 72 | Cross-tissue | Homo sapiens | PMID33184181 | SMC |

Specifically, the mouse T Cell (mTC) dataset was used to illustrate how well each sketching method’s subsample resembles the full dataset (Demo). Four datasets were employed for CTA: human peripheral blood mononuclear cells (PBMC), mouse Brain (mBrain), CELLxGENE minimal (CxG_min), and mouse Aging Cell Atlas (mACA). The Gut and Spleen datasets were used for LT and H, respectively, and all 16 datasets were included in the SMC analysis.

Six of the datasets (those used for CTA, LT, and H) were obtained from links given by their respective literatures, while the remaining 10 were sourced from CELLxGENE (https://cellxgene.cziscience.com/datasets) [2]. PBMC was originally downloaded in Matrix Market format, comprising four files for barcodes, genes, expression values, and metadata; CxG_min was provided in Apache Parquet format; the other 14 datasets were in h5ad format. All datasets had undergone standard quality control by either the original authors or CELLxGENE. Subsequently, we stored and processed all datasets in AnnData format using the Scanpy pipeline [8]. In the experiments, each of the 16 scRNA-seq dataset was normalised with a scale factor of 10,000 and log-transformed (unless already log-normalised in the original files). The top 3,000 highly variable genes (HVGs) were selected if HVGs were not provided or if a different count was not specified in the literature. Lastly, the top 50 principal components (PCs) were computed from these HVGs and used as input for the subsampling methods.

### Overview of scValue

scValue is designed to generate an informative sketch (representative subsample) of large scRNA-seq datasets while preserving critical biological diversity. As illustrated in Fig. 1A, scValue departs from the existing uniform or distance/distribution optimisation-based methods. It first trains a random forest classifier to assign a “data value” to each cell, reflecting how essential it is for distinguishing cell types. The data values then guide subsampling at the cell-type level: a value-weighted allocation determines the target sketch size for each cell type, ensuring that rare yet important cell types are not overlooked; cells with the highest values are selected via a binning procedure by default, producing a smaller dataset that retains the most biologically vital cells.

**Figure 1.**
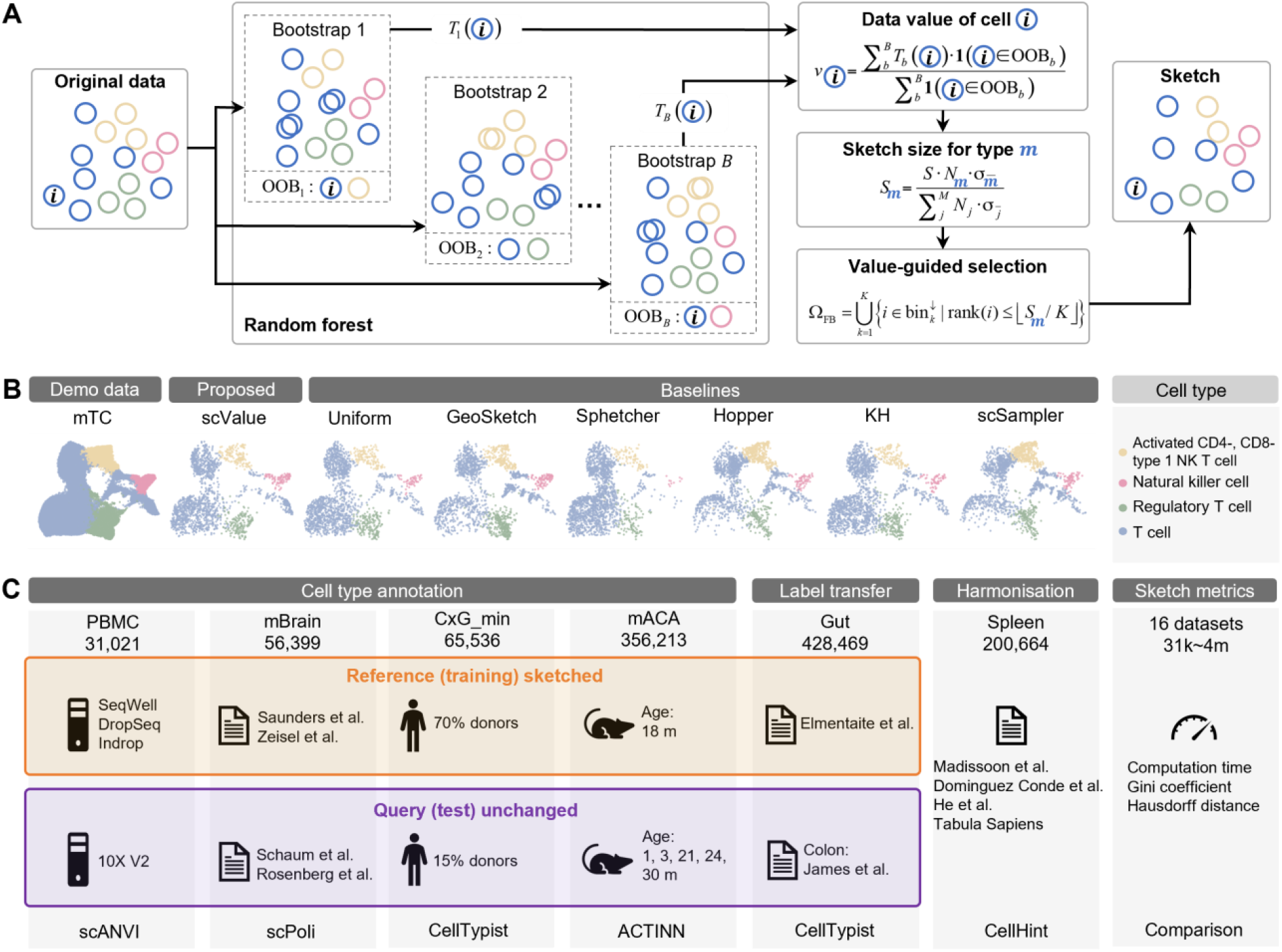
Overview of scValue. (**A**) Given a large scRNA-seq dataset, an informative sketch (representative subsample) is created by scValue through three steps: first, fitting a random forest classifier based on the full dataset with cell-type labels and computing out-of-bag (OOB) estimates as data values of individual cells; second, determining sketch size for each cell type and allocating more representation to cell types with greater value variability; third, dividing the value range into equally sized bins, from each of which top-valued cells are selected to form the resulting subsample. (**B**) Using the mouse T cell (mTC) dataset to demonstrate scValue’s ability of balancing cell-type proportions and enhance cell-type separation, compared to six existing subsampling methods. (**C**) We systematically evaluated scValue against its six counterparts through: four cell-type annotation tasks, each involving a previously studied pair of large scRNA-seq dataset and ML/DL model; a case study of label transfer learning across human gut-colon datasets via CellTypist; another case study of cross-study label harmonisation via CellHint using a human spleen dataset; and finally a sketch metric comparison experiment on 16 large scRNA-seq datasets ranging 31 thousand to four million cells in terms of computation time, Gini coefficient, and Hausdorff distance.

### Data value computation

For a large scRNA-seq dataset containing expression profiles for *N* cells and *M* cell types, we denote the input feature matrix **X** ∈ ℝ ^*N*×*d*^ and the input label vector **Y** ∈{1, 2,…, *M*}^*N*^, respectively. By default, and same as in the other sketching methods, *d* is set as the top number of principal components (PCs) calculated from log-normalised gene counts. Fig. 1A depicts the schematic workflow of scValue using the inputs to create a value-based sketch consisting of *S* cells.

A random forest model [15] comprising *B* decision trees is fitted with (**X, Y**). The *b*-th tree, denoted by *f*_*b*_, is trained on a bootstrap sample drawn with replacement from (**X, Y**). Cell *i* is considered out-of-bag (OOB) if it is not selected in the *b*-th bootstrap, which we denote by *i* ∈ OOB_*b*_. Using only OOB trees to predict for cell *i* and averaging the prediction errors yield the cell’s OOB estimate. This statistic is traditionally used for performance evaluation of random forests [16] and recently revised to facilitate data valuation in machine learning tasks [17]. In our study, each cell’s data value (*v*_*i*_ ∈[0,1]) is computed as the OOB accuracy of cell type prediction

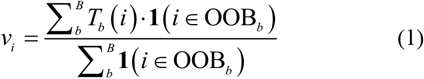

where **1**(*i* ∈ OOB_*b*_) equals 1 if cell *i* is an OOB sample for the *b*-th bootstrap and 0 otherwise; the correctness function *T*_*b*_ (*i*) is given by **1**(*Y*_*i*_ = *f*_*b*_ (**X**_*i*_)), which evaluates to 1 if the *b*-th tree correctly predicts the type of cell *i* and 0 otherwise. Cells of higher value are expected to bring greater benefits for distinguishing cell types when included in the sketch compared to those of lower value.

### Sketch size determination

Sketching is conducted at the cell-type level in a value-weighted manner to improve the cell-type balance in the resulting subset. The number of cells to be subsampled for cell type *m* is determined by considering both the cell type abundance and the standard error of data value

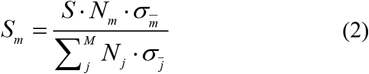

Here, *N*_*m*_ represents the number of cells belonging to type *m* in the original dataset; 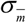 is given by 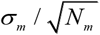, in which *σ* _*m*_ denotes the standard deviation of data value for cells of type *m*. The denominator normalises the value-weighted subsample size across *M* types. In addition, the largest remainder method [18] is employed to ensure that the total allocation for all cell types sums up exactly to.

The rationale behind value-weighted subsampling is as follows. Rare cell types are often more challenging to learn in a random forest due to their underrepresentation, while heterogeneous cell types that contain diverse or complex data also present learning difficulties. Both conditions are expected to result in larger variability in data values. To address this, Eq. (2) seeks to adequately capture rare and heterogenous cell types through the allocation of more cells in the sketch, thereby preserving crucial biological information.

### Value-guided cell selection

Once the sketch size *S*_*m*_ for cell type *m* has been determined, value-guided cell selection can be performed. Let Γ_*m*_ denote the set of all cells of type *m* in the original dataset, and |Γ_*m*_ |= *N*_*m*_. To sketch *S*_*m*_ cells from Γ_*m*_, the full value range [0, 1] is partitioned into *K* bins (by default *K* = 10, i.e., bins of 0.1 intervals). Within the *k*-th bin, let 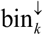 denote the collection of cells ranked by their values *v*_*i*_ in descending order. Uniform bin-wise selection is then performed, picking the top |*S*_*m*_ / *K* | cells from 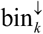:

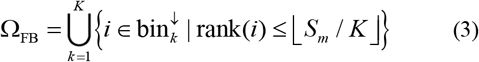

which captures cells across all value ranges to retain a broad snapshot of cell type *m* ‘s variation in value. Hence, we name this strategy as full binning (FB).

Alternatively, two other cell selection strategies can be employed. Mean-threshold binning (MTB) uses the same binning setting as FB; however, only those bins above the mean value are considered. Bins that fall below the mean are combined into a single bin with a lower bound of 0. MTB emphasises cells that exhibit above-average values over less-valuable cells. In contrast, top-pick (TP) directly selects cells with the highest values from each type, focusing on value-maximisation in the final sketch without binning constraints. The three strategies can cater to different analytical needs, achieving varying levels of cell type separation in the sketch. By default, scValue utilises FB since it balances top-valued cell selection and the similarity between the sketch and the original dataset.

### Computational complexity

scValue has *O* (*BdN* log *N*) computational complexity. Here, we review theoretical computational complexities of the six existing subsampling methods, i.e., Uniform (as implemented in the Seurat [7] and Scanpy [8] pipelines), GeoSketch [6], Sphetcher [9], Hopper [10], KH [13], and scSampler [12], as documented in the respective publications. A summary of these complexities, along with that of scValue, is provided in Table 1 for comparison.

Given a large scRNA-seq dataset consisting of *N* cells and *d* features (PCs by default), the sketching objective is to extract cells as a subset that effectively represents the original dataset. Among the seven methods evaluated, five include additional parameters or variables in their computational complexity. Specifically, scValue has a parameter *B*, the number of trees in a random forest, which defaults to 100. TreeHopper, similarly, involves a parameter *B*, denoting the number of partitions of the original dataset, with a default value of 1,000. scSampler also utilises a parameter *B* for the number of partitions, evaluated in its original publication with values of 1 (no partition), 4, and 16 [12]. In the experiments of our study, we set *B* = 4 to balance computational cost and sketch quality. Lastly, KH incorporates a parameter *D*, which denotes the number of random features drawn from the original *d* features.

### Implementation

scValue is implemented in Python. The data value computation step is built upon the *RandomForestClassifier* class (version 1.5.2) of scikit-learn. Parallelisation of tree-fitting is supported to enhance computational speed. By default, scValue accepts the top 50 PCs (*d* = 50) as the latent representation of cells; it can also take gene features or other low-dimensional representations as input. By default, a random forest is constructed with *B* = 100 decision trees. In the value-weighted subsampling step, FB is set as the default cell selection strategy. Furthermore, proportional subsampling can be carried out instead if the original cell types are highly balanced or if users wish to maintain the original cell type proportions.

### Evaluation of scValue

As summarised in Table 1 and illustrated in Fig. 1C, this study evaluated scValue alongside the six existing sketching methods (Uniform, GeoSketch, Sphetcher, Hopper, KH, and scSampler) using the six large scRNA-seq datasets in the ML/DL tasks of cell-type annotation, label transfer, and cell-type harmonisation; then these methods were further assessed with the 16 datasets via sketch metric comparison. In all experiments, each method was applied with its default parameters to generate sketches of each dataset. The tree-partitioning variant of Hopper (TreeHopper) was chosen for its balance between computation time and sketch quality, and scSampler was run with B = 4 partitions to achieve a comparable trade-off. The cell-type annotation tasks were conducted in scenarios where the reference and query data employed consistent cell-type annotation schemes. We replicated four previously studied dataset-model pairs: (1) the human PBMC dataset [19] with the variational autoencoder-based scANVI model [20], originally examined in [21]; (2) the mBrain dataset [22] with the variational autoencoder-based scPoli model [23], investigated in the latter’s study; (3) the CxG_min dataset [24] with the logistic regression-based CellTypist model [25], evaluated in the former’s study; (4) the mACA dataset [26] with the neural networks-based ACTINN model [27], assessed in [28]. For each pair, we used the same dataset partitions (reference and query) described in the corresponding studies. The reference (training) partition was then used to create sketches at 2%, 4%, 6%, 8%, and 10% of the original size via each of the seven subsampling methods, while the query (test) partition remained at its full size. Each resulting sketch was used to train the respective model, which then annotated cell types in the query partition. To ensure robustness, we repeated these steps 10 times with varying random seeds for each sketch, recording the annotation accuracy for each experiment. Moreover, we recorded the accuracy obtained by using the full reference partition for each task (also 10 repeated runs), providing a baseline for assessing the performance drop associated with the different subsampling methods. Further details on partitioning and model training parameters are available in Supplementary Note 1.

The label transfer and harmonisation tasks served as case studies for applying the sketching methods to cross-dataset cell-type transfer learning and cross-dataset cell-type harmonisation, respectively. In the former task, the reference and query data used different annotation conventions. Specifically, we replicated the “large-scale cross-dataset label transfer” tutorial from CellTypist [25], using 10% sketches of the human Gut dataset [29] to annotate a colon query dataset [30] containing 42k cells in full. T-related cells were transferred from each sketch to the query, and we then assessed how closely these transferred annotations matched those obtained when using the full Gut dataset as the reference. For the latter task, the goal was to standardise the cell-type annotation styles across multiple datasets originating from different studies. Following the harmonisation workflow of CellHint [31], we used the human Spleen dataset [25], which encompasses cells from four independent studies. Each subsampling method was applied to sketch 10% of Spleen. CellHint then used these sketches to learn relationships among T cell-related types across the four studies. The performance of cell-type harmonisation was evaluated by comparing the learned relationships from each sketch against those derived from the full Spleen dataset.

Additionally, because most previous studies on existing sketching methods have focused on computation time and Hausdorff distance [6, 9, 10, 12], and the most recent work on scSampler [12] highlights Gini coefficient, we evaluated all three metrics in the sketch metric comparison experiment, using sketches generated by each subsampling method across the 16 datasets (Table 1 and Fig. 1C). Computation time was recorded as the elapsed time (in seconds) to produce each sketch, while Gini coefficient *G* quantified the imbalance in cell-type proportions: lower values indicate a more balanced distribution, and higher values signal greater inequality. This metric is formally defined in [11] as

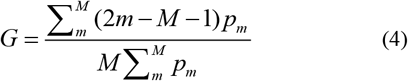

Where *p*_*m*_ represents the proportion of cell type *m* in th full dataset. Consistent with scSampler, we utilised the implementation of Gini coefficient computation available at https://github.com/oliviaguest/gini. Hausdorff distance, denoted by *H*, is defined as the maximum Euclidean distance (*dist*) between any point in the original dataset *X* and the nearest point in *X*_*s*_

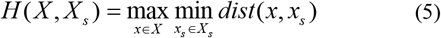

Smaller means that all cells in *X* are well-represented in *X*_*s*_, implying a higher degree of similarity between the original dataset and the subsample. Conversely, larger *H* suggests less similarity.

## Results

### Demonstration: scValue balances cell-type proportions and improves cell-type separation

To illustrate the capacity of scValue to balance cell type proportions while maintaining separation among different cell types, we tested it on the mouse T cell (mTC) dataset [32], comprising ∼90k cells across four cell types. We used *X* scValue and six existing sketching methods to subsample 10% of the original dataset (∼9k cells) and then visualised both the full dataset and each subsample using uniform manifold approximation and projection (UMAP), as shown in Fig. 1B. Overall, scValue, Uniform, and GeoSketch produced sketches that more closely resembled the full distribution compared to Sphetcher, Hopper, KH, and scSampler. Among the former three, both scValue and GeoSketch exhibited better coverage of the relatively rare natural killer cells (pink) by preserving a denser representation of this cell type. Notably, however, scValue achieved clearer separation between T cells (blue) and regulatory T cells (green) than GeoSketch, indicating that scValue not only captures rare populations but also maintains improved resolution between distinct cell types. These observations hence underscore the ability of scValue to effectively balance cell type proportions and enhance separation among cell types, with the potential of benefiting ML/DL tasks.

### Evaluation: scValue outperforms existing sketching methods in ML/DL-based cell-type annotation tasks

To assess how well scValue preserves essential information in sketches for ML/DL-based cell type annotation, we evaluated the method against its six counterparts on four dataset-model pairs: PBMC with scANVI [21], mBrain with scPoli [23], CxG_min with CellTypist [24], mACA with ACTINN [28]. Each dataset was split into a reference set (for training) and a full-sized query set (for validation). We then generated sketches of the reference set at varying fractions (2-10% of the original dataset) using the seven subsampling methods. Subsequently, the trained models were used to infer cell types on the full query set; and this procedure was repeated ten times to ensure robust performance estimates. Fig. 2 presents the annotation accuracies in boxplots and Tables S1–S4 summarise the average and standard deviation of accuracies for each sketching method at each subsampling percentage in each dataset-model pair’s experiment.

**Figure 2.**
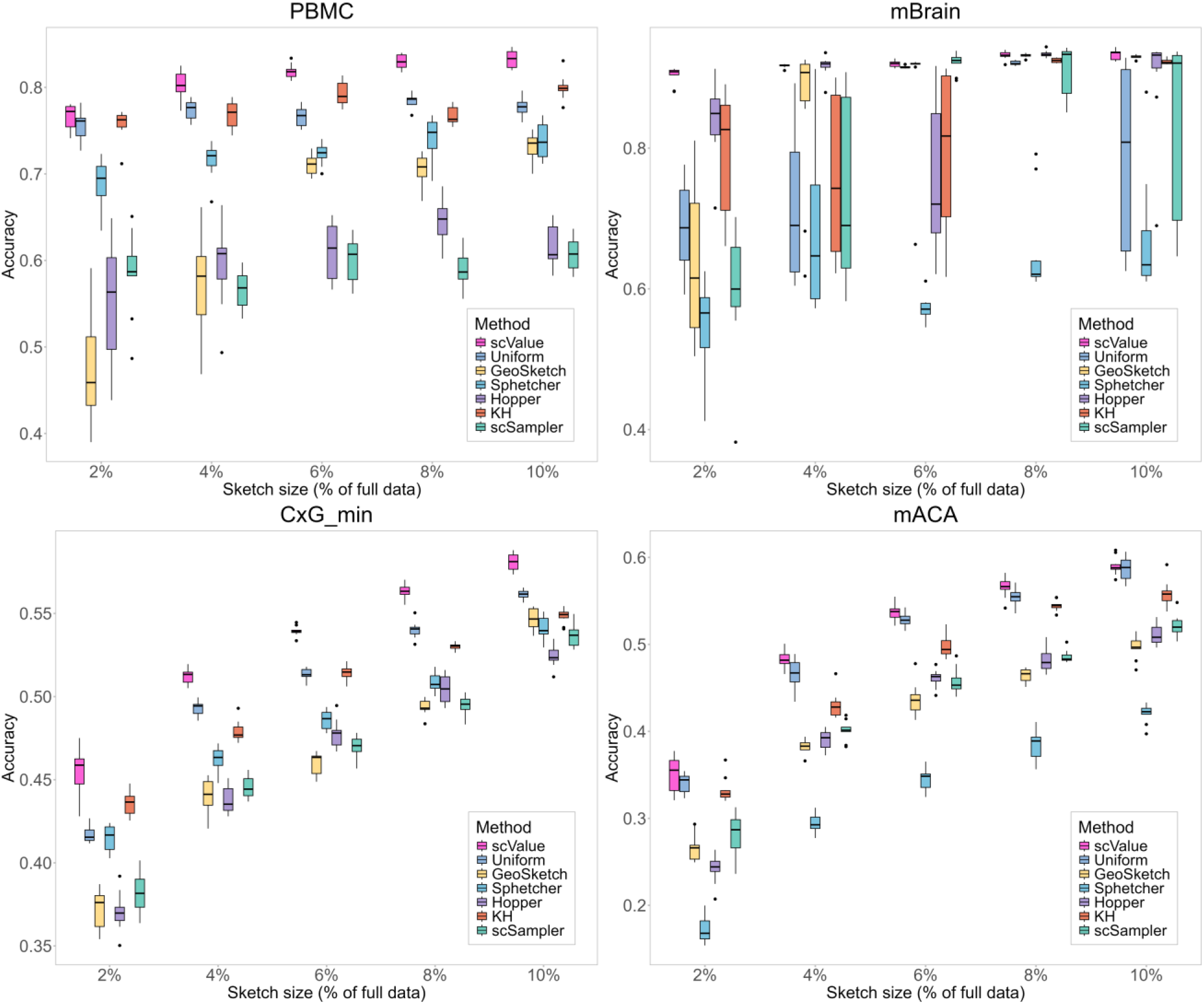
scValue outperforms existing sketching methods in ML/DL-based cell-type annotation tasks. We evaluated scValue against Uniform, GeoSketch, Sphetcher, Hopper, KH, and scSampler on four previously studied dataset-model pairs: PBMC with variational autoencoder-based scANVI, mBrain with variational autoencoder-based scPoli, CxG_min with logistic regression-based CellTypist, mACA with neural network-based ACTINN. For each dataset, sketches of the reference partition were created at varying fractions (2-10%) and then used to train the corresponding model for annotating the full query partition.

For PBMC, scValue consistently outperformed the other methods across all sketch sizes. This was especially evident at smaller sketch percentages (2% and 4%), where accuracy values tended to be more volatile. Indicated by the narrower interquartile ranges in the boxplots, performance generally stabilised as the sketch size increased. Notably, only scValue and Sphetcher exhibited the desirable trend of steadily increasing annotation accuracy with larger sketch sizes. At 10%, scValue reached an average accuracy of 0.8330±0.0103, close to the 0.8635±0.0050 achieved using the full reference (Table S1). In contrast, no other method surpassed an average accuracy of 0.8 at 10%, underscoring scValue’s superior performance.

In the learning task on mBrain, although several methods approached or even slightly surpassed the full-reference accuracy (0.9320±0.0047) at higher sketch percentages, their performance was considerably less stable than scValue’s. In the challenging regime of 2-4% sketches, scValue maintained consistently high accuracy while other methods varied more widely. For instance, at 2%, scValue (0.9033±0.0122) surpassed all other methods by a substantial margin (Table S2). As the sketch size increased, counterparts such as Uniform, Hopper, or scSampler occasionally matched or exceeded scValue’s accuracy for specific percentages (e.g., 6% or 8%), but they did so with greater variability across repeated runs. Importantly, scValue was the only method that maintained an upward trend in accuracy as the sketch size increased, reflecting its robustness to small data and ability to leverage larger training subsets. Interestingly, Hopper achieved a slightly higher accuracy than the full reference at 8% (0.9338 vs. 0.9320), suggesting that mBrain is relatively easy to learn for certain methods, although this improvement was marginal.

For CxG_min, all methods improved in accuracy as the sketch size grew, but scValue again outperformed the others across the entire range. Although its boxplots occasionally exhibited longer whiskers (indicating certain degrees of variability across repeated runs), scValue’s lowest accuracy within each whisker was still higher than the highest accuracy of any competing method in the same sketch-size group (especially notable at 4-10%). At 10%, scValue reached an average accuracy of 0.5811±0.0052, closer to the full-data accuracy of 0.6847±0.0045 than any other method (the next best, Uniform, achieved 0.5615±0.0027; Table S3). These results suggest that scValue retains a more representative subset of the data, ensuring better performance even for relatively complex datasets.

mACA was the most challenging dataset, with overall accuracies ranging from ∼0.2 to ∼0.6. This difficulty likely arose from the larger number of cell types (197), creating a more complex annotation task. Despite this challenge, scValue again led for the top performance at all sketch sizes, highlighting its resilience in small-data scenarios. At 10%, scValue achieved an average accuracy of 0.5901±0.0098, surpassing Uniform (0.5871±0.0129) and substantially exceeding methods like GeoSketch, Sphetcher, or scSampler (Table S4). The gap of 0.1275 from the 0.7176±0.0046 accuracy obtained using the full reference was larger than that observed in PBMC (0.0305), mBrain (−0.0018), and CxG_min (0.1036). We speculate that the higher complexity of mACA, with 197 distinct cell types, imposed a greater challenge for any subsampling approach to produce a near-complete representation of the cellular landscape. Nonetheless, mACA still demonstrated stable learning trends overall: all methods gradually improved as more data were included in the sketches.

Across the four dataset-model pairs, scValue achieved the highest (or occasionally near-highest) annotation accuracies, frequently outperforming alternative sketching methods by notable margins, particularly at smaller sketch sizes where data scarcity can significantly degrade model performance. Importantly, at the 10% sketch level, scValue’s average accuracies were closest to those derived from the full reference dataset, highlighting its effectiveness in retaining critical information needed for robust cell type annotation. Overall, these findings confirm that scValue strikes a favourable balance between efficiency (data reduction) and model performance, making it a strong choice for sketching large single-cell datasets in ML/DL-based cell type annotation tasks.

### Case study: scValue enables improved CellTypist label transfer learning

To examine how scValue performs when reference and query data have divergent annotation styles, we conducted a label transfer learning experiment using the Gut (428k cells) [29] and immune (42k cells) [30] datasets from the CellTypist tutorial [25]. Following the tutorial’s workflow, we first reduced the 428k Gut dataset to 55k balanced cells (ensuring each cell type was equally represented). Because the dataset was already well-balanced, we used proportional subsampling rather than value-weighted subsampling and then applied scValue to produce a 10% sketch, while the immune dataset (42k cells) remained unchanged. We trained a CellTypist model [24] on each 10% sketch (generated by scValue and the six baseline subsampling methods) and then transferred its annotations to selected T-cell populations (“Activated CD4 T,” “Th1,” “Tfh,” “CD8 T,” “cycling gd T,” “Tcm,” “gd T,” “Th17,” and “Treg”) in the immune query dataset. To evaluate performance, the transferred annotations were compared against the query dataset’s original annotations. Fig. 3 shows dot plots contrasting these predictions with the original alignment obtained using the full reference data in the tutorial.

**Figure 3.**
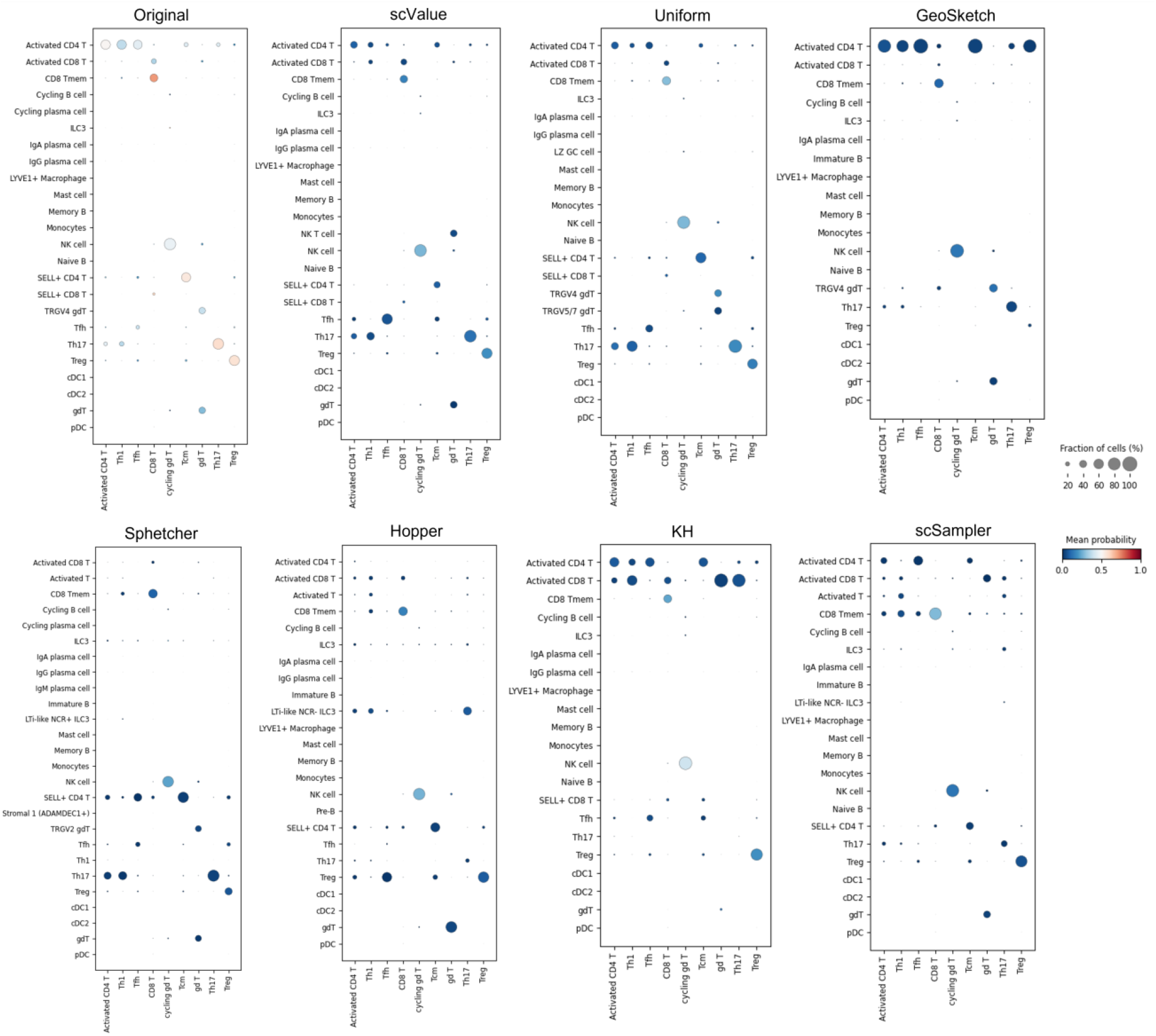
scValue enables improved CellTypist label transfer learning. In the case study of employing CellTypist to transfer labels from the Gut (reference) to the Colon (query) dataset, scValue’s sketch preserved T-cell labels most similar to the full reference and even surpassed the latter in the “Tfh” cell type, as reflected in the larger fraction of cells being correctly predicted for this type.

Several trends can be observed from the plots. Among all methods, scValue and Uniform yielded label transfers that most resembled the original ones. In addition, scValue not only preserved but also improved the correct classification of “Tfh” cells, indicating its strength in retaining subtle yet important features of cell types. The other five sketching methods showed higher levels of different assignments:

- GeoSketch had a larger proportion of “CD8 T” cells and “Treg” cells classified as “Activated CD4 T”.
- Sphetcher did not predict any “Activated CD4 T cells”.
- Hopper inferred the majority of “Activated CD4 T” cells as other T subtypes.
- KH displayed a noticeable tendency to label “Activated CD4 T” as “Activated CD8 T”.
- scSampler classified a large portion of “Activated CD4 T” as “CD8 Tmem”, and some “CD8 T” as “SELL+ CD4 T”.

Subsampling the reference data naturally led to lower label transfer probabilities, as visually reflected by more blue bubbles (indicating reduced confidence) in each sketch’s dot plot. Nevertheless, scValue emerged as a top-performing subsampling strategy, preserving enough critical information to sustain accurate label transfer.

### Case study: scValue facilitates improved label harmonisation with CellHint

To explore how effectively scValue captures complex cell-type relationships when integrating scRNA-seq data from multiple sources with divergent annotation styles, we followed the CellHint tutorial workflow [31] to harmonise T-cell annotations across four independent studies, collectively comprising 200k cells in the Spleen dataset [25]. Each study provides a slightly different nomenclature for T-cell subtypes, making robust cross-study alignment crucial for constructing a standardised cell atlas.

In this experiment, we first generated a 10% sketch of the Spleen dataset (i.e., ∼20k cells) using scValue. Notably, because the original dataset already contained various distinct T-cell labels from multiple sources, our goal was to preserve these fine-grained subpopulations in the sketch. We then applied CellHint [31] to learn hierarchical relationships among the T-cell annotations from the four studies, using the scValue-derived sketch as input. For comparative evaluation, the other six sketching methods also created 10% subsets, which were fed through the same pipeline. Fig. 4 depicts the resulting tree plots representing inter-study label relationships, juxtaposed with the original tree constructed from the full 200k dataset.

**Figure 4.**
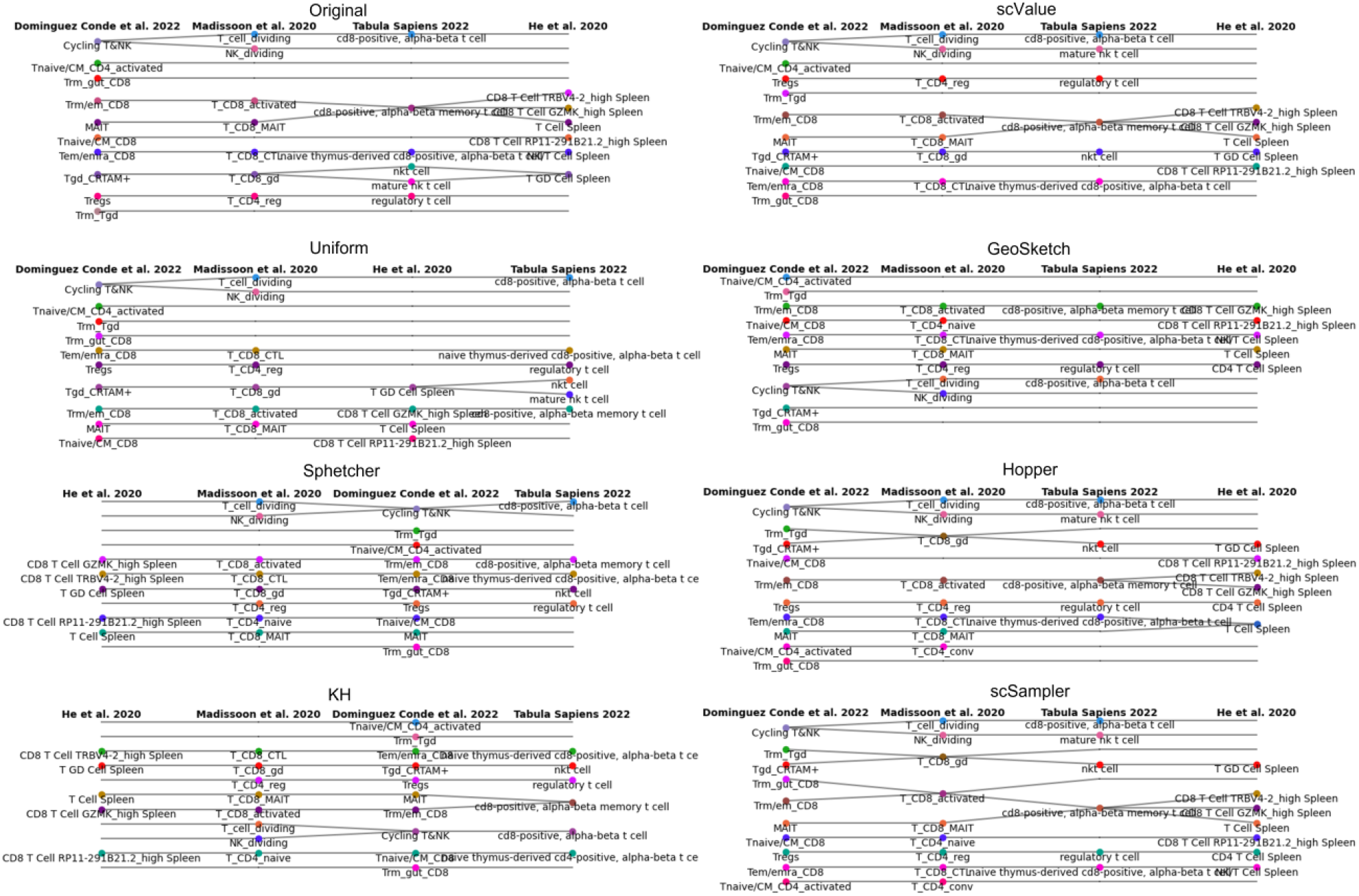
scValue facilitates improved label harmonisation with CellHint. In the case study of label harmonisation for the multi-study Spleen dataset via CellHint, scValue’s sketch reproduced inter-study T-cell subtype relationships more closely to those obtained with the full data than sketches of other subsampling methods.

Of all methods tested, scValue performed best in reproducing the fine-grained relationships among T-cell subtypes observed in the full dataset’s tree. This reflects scValue’s ability to maintain critical cellular diversity in reduced subsets, even under conditions where the nomenclature varies across studies. Other methods exhibited diverse degrees of different matching compared to the original tree, including deletion of existing T-cell subtype relationships and/or introductions of new ones:

- Uniform resulted in a reordering of studies and removed relationships involving the “CD8-positive, alpha-beta memory T cell” and “T Cell Spleen”.
- GeoSketch related “Tnaive/CM CD8” to “T_CD4 naïve”.
- Sphetcher both altered the study order and linked “Tnaive/CM CD8” to “T_CD4 naïve”.
- Hopper merged “Trm_Tgd” and “Tgd_CRTAM+” into “T_CD8_gd”.
- KH similarly led to a change of study ordering and related “Tnaive/CM CD8” to “T_CD4 naïve”.
- scSampler introduced several additional relationships including linking “Tnaive/CM_CD4_activated” with “T_CD4_conv”, merging “Trm_Tgd” and “Tgd_CRTAM+” into “T_CD8_gd”, and integrating “Trm_gut_CD8” and “Trm/em_CD8” into “T_CD8_activated”.

These inconsistencies likely stem from each method’s differing priorities in selecting representative cells. Approaches that under-sample or over-sample particular subpopulations can distort the underlying biological relationships, particularly in datasets with nuanced cell-type definitions that vary by study. In contrast, scValue’s built-in balancing and value-based selection strategies appear to safeguard these subtle distinctions.

### Comparison: scValue yields efficient computation time, best Gini coefficient, and uniform-like Hausdorff distance

To conclude the evaluations, we benchmarked scValue against the six other methods using the 16 datasets of varying size (Table 1), focusing on computation time, Gini coefficient, and Hausdorff distance. For each dataset, a 10% sketch was generated by each method. The scatter plots in Fig. 5 illustrate computation time (x-axis) versus Gini coefficient (y-axis), with bubble size indicating the sketch’s Hausdorff distance from the full dataset. The detailed statistics for producing the plots are provided in Table S5. A major observation is that scValue (shown in pink) typically clustered toward the lower ends of both axes, reflecting its short runtime and low Gini coefficient (balanced cell-type proportions). Meanwhile, the moderate bubble size indicates that scValue’s sketch remains reasonably close to the original data distribution. It should be noticed that, for the mEmbryo dataset (containing ∼3.2 million cells), Sphetcher and KH did not complete their runs and thus no sketches were included in the relevant plots; similarly, for the Fetal dataset (about four million cells), Hopper, Sphetcher, and KH did not finish running and were therefore excluded from the results.

**Figure 5.**
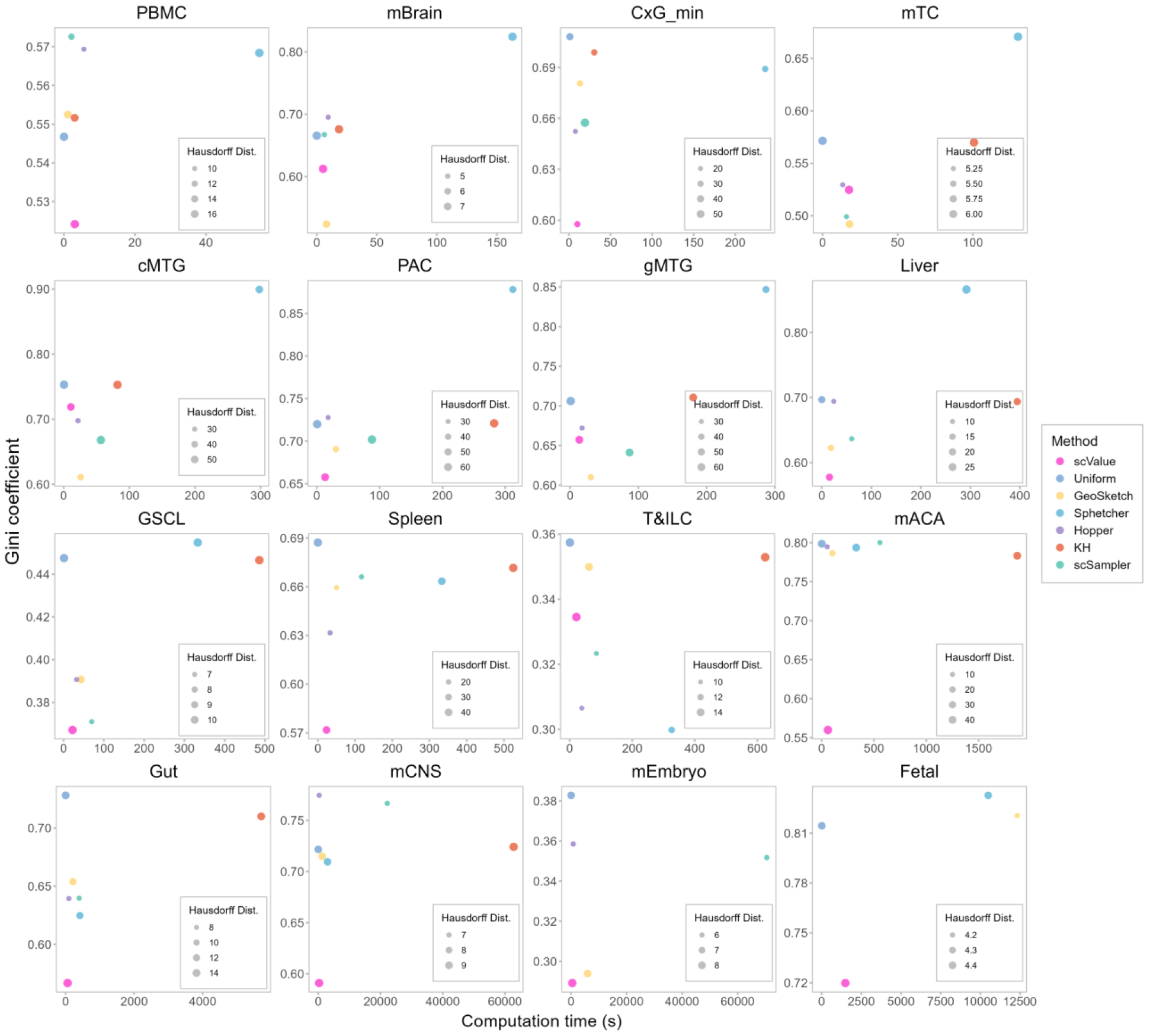
scValue yields efficient computation time, best Gini coefficient, and uniform-like Hausdorff distance. We evaluated scValue against its counterparts in terms of three sketch quality metrics using the 16 datasets containing between 31 thousand and four million cells, spanning four to 197 cell types and including samples from more than 10 tissues across four species.

From Table 3, which ranks each method by its average performance on the three metrics across the datasets, scValue stood out for its computation time rank of 2.4±0.8, second only to the simple Uniform approach, and best Gini coefficient rank of 2.1±1.5. The Hausdorff Distance rank of 5.3±1.1 was slightly lower than Uniform (5.5±1.2) and KH (5.4±1.3), confirming that scValue’s sketches did not deviate excessively from the original distribution. These results suggest that scValue strikes a distinct trade-off: it excels at capturing the full diversity of cell populations and is computationally efficient, yet it can exhibit a slightly higher distributional shift compared to alternative methods that are based on distance-optimisation (especially, GeoSketch, Hopper, and scSampler). Nevertheless, this shift remains comparable or better than Uniform’s performance, reinforcing that scValue offers a robust balance between cell-type representativeness, efficiency, and data fidelity. Finally, Fig. S1 further illustrates how scValue’s computational time is nearly independent of the sketch size (tested from 2% to 10%) using the Spleen dataset as an example, which resonates with *O* (*BdN* log *N*) complexity. This constant-time characteristic emphasises the practical scalability of scValue, which is an important consideration for large-scale single-cell applications where speed matters.

## Discussion

In this study, we introduced scValue, a novel subsampling method devised for large-scale single-cell transcriptomic data. Conceptually different from existing approaches that optimise distance (e.g., GeoSketch, Sphetcher, Hopper, and scSampler) or distribution (e.g., KH) metrics, scValue prioritises each cell by a data value that reflects its utility in distinguishing cell types (Fig. 1A). Specifically, given a large scRNA-seq dataset, scValue begins with data value computation: a random forest classifier is fit to the full dataset using cell-type labels; and each cell’s OOB estimate becomes its data value, capturing how strongly that cell contributes to classification. Next, a target sketch size per cell type is determined by accounting for both abundance and the variability in data values across that cell type; cell types with greater variance in data values receive proportionally more cells in the sketch, ensuring that rare or biologically intricate cell types are sufficiently represented. Lastly, within each cell type, scValue applies the full binning (FB) strategy to select cells at different value levels: the value range is divided into equally sized bins, from which the top-valued cells are included, thereby preserving original data diversity while still favouring cells with higher classification utility. Through a demonstration on the mTC dataset, scValue is shown to balance cell-type proportions and enhance cell-type separation in the UMAP plot (Fig. 1B).

We then systematically compared scValue against six subsampling methods (Uniform, GeoSketch, Sphetcher, Hopper, KH, and scSampler) in a series of machine/deep learning tasks (Fig. 1C). In the cell type annotation tasks (Fig. 2), these methods were evaluated with four previously studied dataset-model pairs, encompassing 31 to 356 thousand cells (PBMC, mBrain, CxG_min, and mACA) and a variety of models (variational autoencoder-based scANVI and scPoli, logistic regression-based CellTypist, and neural network-based ACTINN). scValue consistently achieved the highest or near-highest annotation accuracies, maintained stable performance across repeated runs, and improved accuracies steadily as sketch size increased. Notably, at a 10% sketch level, scValue’s annotation results were closest to those trained on the full reference set. In the case study of CellTypist label transfer from the Gut (reference) to the Colon (query) dataset (Fig. 3), scValue’s sketch preserved T-cell labels most similar to the full reference and even surpassed the latter in the “Tfh” cell type. Moreover, in the case study of CellHint label harmonisation for the multi-study Spleen dataset (Fig. 4), scValue’s sketch reproduced inter-study T-cell subtype relationships more closely to those obtained with the full data than sketches of other subsampling methods. Finally, as depicted in Fig. 5 and Table 3, across a broader collection of 16 scRNA-seq datasets spanning 31 thousand to four million cells (Table 1), scValue outperformed or matched its counterparts in terms of computation Time (second only to simple Uniform, thus highly scalable for large datasets), Gini coefficient (top ranked, indicating well-balanced cell-type representations), and Hausdorff distance (comparable to Uniform, implying no excessive distributional deviation from the original datasets).

Overall, scValue strikes a favourable balance between computational efficiency, preservation of critical biological information, and reliable performance in downstream machine/deep learning (ML/DL) tasks. By using OOB estimates from a random forest [17], scValue introduces a biologically interpretable “cell importance” measure, i.e., a higher data value signals a cell more characteristic of its type. This represents a departure from the purely geometric or density-based criteria that underlie existing approaches [6, 9, 10, 12, 13], linking the subsampling procedure directly to supervised learning performance. In addition to its data value computation step, scValue’s value-weighted sketch size determination and value-guided cell selection go beyond simple proportional sampling; together, these elements ensure that rare or heterogeneous cell populations receive adequate representation, mitigating the risk of overshadowing by more abundant cell types. Indeed, the method’s emphasis on balancing cell-type representation and prioritising high-valued cells underpins its superior performance in cell type annotation, transfer learning, and harmonisation tasks. An intriguing observation, however, is that top performance does not strictly require maximising both “balance” and “value selection” simultaneously. For example, in the mBrain-scPoli experiment, scValue did not show the top Gini coefficient (Fig. 5) yet achieved the highest accuracies at most sketch sizes (Fig. 2 and Table S2). This highlights the need for future experiments to dissect the relative contributions of balanced cell-type proportions versus top-valued cell selection and whether one factor can outweigh the other, toward robust ML/DL outcomes.

Regarding the computation time observed in the sketch metric comparison (Fig. 5 and Table 3), scValue’s practical efficiency can be explained as follows. Since the target sketch size *S* is typically considerably larger than log(*N*) for typical sketching applications in single-cell analysis [6], it can be discerned from Table 2 that the ranking of theoretical computational complexity in ascending order is: Uniform < GeoSketch < scValue < (Tree)Hopper ≈ scSampler (with partitioning) < Sphetcher < KH. However, actual computation time may deviate due to several factors [33], such as the constant and lower-order terms disregarded in Big-O notation, dependencies on specific characteristics of the data, memory management overhead, among others. Moreover, as a random forest in scValue can be easily parallelised, this method may therefore achieve faster or comparable runtimes to the other sketching methods (including GeoSketch) in practice, particularly for large datasets, as illustrated in the results.

**Table 2.** Theoretical computational complexities for the seven sketching methods.

| Method | Theoretical complexity |
| --- | --- |
| scValue | $O(BdN \log N)$ with $B$ trees and $d$ features |
| Uniform | $O(1)$ |
| Geosketch | $O(dN \log N)$ with $d$ features |
| Hopper | $O(NS)$ |
| | $O\left(\frac{NS}{B}\right)$ for TreeHopper with $B$ partitions |
| Sphetcher | $O(\sum_i \vert L_i\vert )$ for each iteration until $L$ converges, where $\vert L_i\vert $ is the size of the $i$-th spherical subset |
| KH | $O(dDNS)$ with $d$ features and $D$ random features |
| scSampler | $O(NS)$ |
| | $O\left(\frac{NS}{B}\right)$ if split by $B$ subsets |

**Table 3.** Rankings of each subsampling method by its average performance on the three metrics across the 16 datasets.

| Method | Computation time rank | Gini coefficient rank | Hausdorff distance rank |
| --- | --- | --- | --- |
| scValue | $2.4 \pm 0.8$ | $2.1 \pm 1.5$ | $5.3 \pm 1.1$ |
| Uniform | $1.0 \pm 0.0$ | $5.4 \pm 1.6$ | $5.5 \pm 1.2$ |
| GeoSketch | $3.9 \pm 0.5$ | $2.6 \pm 1.4$ | $2.7 \pm 0.6$ |
| Sphetcher | $6.1 \pm 1.0$ | $5.1 \pm 2.3$ | $4.7 \pm 1.6$ |
| Hopper | $3.1 \pm 0.9$ | $4.0 \pm 1.5$ | $1.3 \pm 0.5$ |
| KH | $6.5 \pm 0.8$ | $5.0 \pm 1.3$ | $5.4 \pm 1.3$ |
| scSampler | $4.9 \pm 0.9$ | $3.7 \pm 1.7$ | $2.6 \pm 1.9$ |

Despite these strengths, several challenges remain. First, experimental results suggest that the number of cell types directly influences the performance gap between scValue’s sketches and the full data. Specifically, datasets with many cell types like CxG_min (164 cell types) and mACA (197 cell types) showed noticeably higher performance gaps (0.1036 and 0.1275, respectively) than PBMC (0.0305) and mBrain (−0.0018), both of which contain only 10 cell types. It is not surprising that as the number of cell types increases, the complexity and heterogeneity of the data also escalate, making it more challenging to obtain representative subsamples by scValue to capture all biological signals (though the method still outperformed its counterparts in almost all cases). Consequently, refining the number of cell types in the original data appears necessary to achieve performance comparable to that of the full dataset. Second, from an algorithmic standpoint, scValue’s reliance on accurately learned data values assumes reasonably clean gene expression data. Noisy datasets with duplicate or near-duplicate cells can inflate OOB estimates for artefactual “redundant” cells, confusing scValue’s interpretation of which populations are rarer and which cells are truly high-value. Careful quality control thus becomes essential for ensuring that scValue appropriately assigns high-value scores and maintains sketch quality. Third, another limitation arises in distribution-sensitive bioinformatics tasks (e.g., differential expression analysis and gene regulatory network inference [6]), where scValue’s sketches that are approximately uniform might still be inadequate. Caution is therefore advised before applying scValue outside of ML/DL contexts, where subtle distributional details are crucial. Fourth, a general caveat in scRNA-seq ML/DL workflows is that after subsampling, the number of cells may become comparable to or even less than the number of genes, amplifying dimensionality issues [34]. It remains critical to use an appropriate number of HVGs or suitably designed model architectures to mitigate the curse of dimensionality.

To make a final point, scValue can be envisioned as a generic data valuation and subsampling framework for single-cell data. Although the current method focuses on cell-type labels, future extensions could incorporate other meta-information, e.g., developmental timing, disease status, tissue source, and species, when utilising such information to build predictive models at scale. Besides, subsampling may be reframed as removing low-value cells to identify a minimal dataset yielding near-optimal predictive performance. Such a “minimal-viable sketch” concept could be highly relevant for studies constrained by computing resources yet eager to preserve salient biological signals in ML/DL applications.

### Key Points

- Value-based subsampling of large scRNA-seq data tied to classification utility: scValue introduces a novel “value-based” strategy that uses out-of-bag (OOB) estimates from a random forest to quantify each cell’s importance (value) for distinguishing its cell type. By linking subsampling directly to classification performance, scValue targets the most informative cells for efficient downstream ML/DL tasks.
- Balanced but biologically rich sketches: Through variance-guided allocation, scValue ensures that rare or complex cell types receive proportionally more top-valued cells in the sketch, preserving critical biological information.
- Consistently high ML/DL accuracy: In benchmarks spanning cell-type annotation, cross-dataset label transfer, and label harmonisation, scValue outperforms (or occasionally matches) existing subsampling methods. Notably, it retains model performance close to that of training on the full dataset.
- Efficiency and scalability: scValue runs quickly on large scRNA-seq datasets (tens of thousands to millions of cells), creating balanced sketches with near-uniform data distributions.

## Acknowledgements

We thank the high-throughput sequencing and high-performance computing platform of the Institute of Systems Medicine for technical support.

## Conflict of Interest

none declared.

## Funding

This work has been supported by National Natural Science Foundation of China (32300560); CAMS Innovation Fund for Medical Sciences (CIFMS) [2021-I2M-1-061, 2022-I2M-2-004, 2023-I2M-2-005]; Non-profit Central Research Institute Fund of Chinese Academy of Medical Sciences [2022-RC416-01]; NCTIB Fund for R&D Platform for Cell and Gene Therapy, the Suzhou Municipal Key Laboratory [SZS2022005]; the Special Research Fund for Central Universities, Peking Union Medical College (3332024089). Chinese Academy of Medical Sciences & Peking Union Medical College, Union Medical College Young Scholar Support Program, No. 2023086; China Postdoctoral Science Foundation, the Postdoctoral Fellowship Program (Crade B), Grant Number GZB20230084.

## Data availability

PBMC is accessible in the Matrix Market format from https://portals.broadinstitute.org/single_cell/study/SCP424/single-cell-comparisonpbmc-data. mBrain is acquired by downloading the mouse_brain_normalized.h5ad file from the dataset directory at https://github.com/theislab/scArches-reproducibility. CxG_min is obtained from https://github.com/theislab/scTab/tree/devel, where the minimal subset of the training and test data were used in our study. mTC is available under “Major cell cluster: T cells” at https://cellxgene.cziscience.com/collections/45d5d2c3-bc28-4814-aed6-0bb6f0e11c82. cMTG is collected under “Chimpanzee: Great apes study” from https://cellxgene.cziscience.com/collections/4dca242c-d302-4dba-a68f-4c61e7bad553. PAC is obtained under “Dissection: Primary auditory cortex(A1)” from https://cellxgene.cziscience.com/collections/d17249d2-0e6e-4500-abb8-e6c93fa1ac6f. gMTG is downloaded under “Gorilla: Great apes study” from https://cellxgene.cziscience.com/collections/4dca242c-d302-4dba-a68f-4c61e7bad553. Liver is available under “All cells from human liver dataset” at https://cellxgene.cziscience.com/collections/74e10dc4-cbb2-4605-a189-8a1cd8e44d8c. GSCL is sourced under “Human Somatic Cell Lineage” from https://cellxgene.cziscience.com/collections/661a402a-2a5a-4c71-9b05-b346c57bc451. Spleen is downloaded from https://celltypist.cog.sanger.ac.uk/Resources/Organ_atlas/Spleen/Spleen.h5ad. T&ILC is collected under “T & innate lymphoid cells” from https://cellxgene.cziscience.com/collections/62ef75e4-cbea-454e-a0ce-998ec40223d3. mACA is sourced from https://figshare.com/articles/dataset/Processed_files_to_use_with_scanpy_/8273102/3; we used the tabula-muris-senis-bbknn-processed-official-annotations.h5ad file (the Version 3 atlas). Gut is obtained from https://cellgeni.cog.sanger.ac.uk/gutcellatlas/Full_obj_raw_counts_nosoupx.h5ad. mCNS is collected under “Major cell cluster: CNS neurons” from https://cellxgene.cziscience.com/collections/45d5d2c3-bc28-4814-aed6-0bb6f0e11c82. mEmbryo is available under “Major cell cluster: Mesoderm” at https://cellxgene.cziscience.com/collections/45d5d2c3-bc28-4814-aed6-0bb6f0e11c82. Fetal is sourced under “Survey of human embryonic development” from https://cellxgene.cziscience.com/collections/c114c20f-1ef4-49a5-9c2e-d965787fb90c.

## Notes

### Competing Interest Statement

The authors have declared no competing interest.

### Summary of Updates

Article's content reorganised; figures and tables revised; two additional cell-type annotation experiments added; discussion added.

## References

1. Regev A, Teichmann SA, Lander ES et al. The human cell atlas, elife 2017;6:e27041.

2. CZI Single-Cell Biology Program, Abdulla S, Aevermann B et al. CZ CELL× GENE Discover: A single-cell data platform for scalable exploration, analysis and modeling of aggregated data, BioRxiv 2023:2023.2010. 2030.563174.

3. Deng Y, Chen P, Xiao J et al. SCAR: single-cell and Spatially-resolved Cancer Resources, Nucleic Acids Research 2024;52:D1407–D1417.

4. Deng Y, Lu Y, Li M et al. SCAN: Spatiotemporal Cloud Atlas for Neural cells, Nucleic Acids Research 2024;52:D998–D1009.

5. Johnston KG, Grieco SF, Nie Q et al. Small data methods in omics: the power of one, Nature Methods 2024:1–6.

6. Hie B, Cho H, DeMeo B et al. Geometric Sketching Compactly Summarizes the Single-Cell Transcriptomic Landscape, Cell Syst 2019;8:483–493 e487.

7. Satija R, Farrell JA, Gennert D et al. Spatial reconstruction of single-cell gene expression data, Nat Biotechnol 2015;33:495–502.

8. Wolf FA, Angerer P, Theis FJ. SCANPY: large-scale single-cell gene expression data analysis, Genome Biol 2018;19:15.

9. Do VH, Elbassioni K, Canzar S. Sphetcher: Spherical Thresholding Improves Sketching of Single-Cell Transcriptomic Heterogeneity, iScience 2020;23:101126.

10. DeMeo B, Berger B. Hopper: a mathematically optimal algorithm for sketching biological data, Bioinformatics 2020;36:i236–i241.

11. Johnson ME, Moore LM, Ylvisaker D. Minimax and maximin distance designs, Journal of statistical planning and inference 1990;26:131–148.

12. Song D, Xi NM, Li JJ et al. scSampler: fast diversity-preserving subsampling of large-scale single-cell transcriptomic data, Bioinformatics 2022;38:3126–3127.

13. Baskaran VA, Ranek J, Shan S et al. Distribution-based sketching of single-cell samples. In: Proceedings of the 13th ACM International Conference on Bioinformatics, Computational Biology and Health Informatics. 2022, p. 1–10.

14. Ma Q, Xu D. Deep learning shapes single-cell data analysis, Nat Rev Mol Cell Biol 2022;23:303–304.

15. Breiman L. Random forests, Machine learning 2001;45:5–32.

16. Efron B. Jackknife-after-bootstrap standard errors and influence functions, Journal of the Royal Statistical Society Series B: Statistical Methodology 1992;54:83–111.

17. Kwon Y, Zou J. Data-oob: Out-of-bag estimate as a simple and efficient data value. In: International Conference on Machine Learning. 2023, p. 18135–18152. PMLR.

18. Balinski ML, Young HP. Fair representation: meeting the ideal of one man, one vote. Rowman & Littlefield, 2010.

19. Abdelaal T, Michielsen L, Cats D et al. A comparison of automatic cell identification methods for single-cell RNA sequencing data, Genome Biol 2019;20:194.

20. Xu C, Lopez R, Mehlman E et al. Probabilistic harmonization and annotation of single-cell transcriptomics data with deep generative models, Mol Syst Biol 2021;17:e9620.

21. Du ZH, Hu WL, Li JQ et al. scPML: pathway-based multi-view learning for cell type annotation from single-cell RNA-seq data, Commun Biol 2023;6:1268.

22. Lotfollahi M, Naghipourfar M, Luecken MD et al. Mapping single-cell data to reference atlases by transfer learning, Nat Biotechnol 2022;40:121–130.

23. De Donno C, Hediyeh-Zadeh S, Moinfar AA et al. Population-level integration of single-cell datasets enables multi-scale analysis across samples, Nature Methods 2023;20:1683–1692.

24. Fischer F, Fischer DS, Mukhin R et al. scTab: scaling cross-tissue single-cell annotation models, Nature Communications 2024;15:6611.

25. Dominguez Conde C, Xu C, Jarvis LB et al. Cross-tissue immune cell analysis reveals tissue-specific features in humans, Science 2022;376:eabl5197.

26. Tabula Muriss C. A single-cell transcriptomic atlas characterizes ageing tissues in the mouse, Nature 2020;583:590–595.

27. Ma F, Pellegrini M. ACTINN: automated identification of cell types in single cell RNA sequencing, Bioinformatics 2020;36:533–538.

28. Chen J, Xu H, Tao W et al. Transformer for one stop interpretable cell type annotation, Nat Commun 2023;14:223.

29. Elmentaite R, Kumasaka N, Roberts K et al. Cells of the human intestinal tract mapped across space and time, Nature 2021;597:250–255.

30. James KR, Gomes T, Elmentaite R et al. Distinct microbial and immune niches of the human colon, Nat Immunol 2020;21:343–353.

31. Xu C, Prete M, Webb S et al. Automatic cell-type harmonization and integration across Human Cell Atlas datasets, Cell 2023;186:5876–5891 e5820.

32. Qiu C, Martin BK, Welsh IC et al. A single-cell time-lapse of mouse prenatal development from gastrula to birth, Nature 2024;626:1084–1093.

33. Cormen TH, Leiserson CE, Rivest RL et al. Introduction to algorithms. Cambridge, Massachusetts: MIT press, 2009.

34. Altman N, Krzywinski M. The curse (s) of dimensionality, Nat Methods 2018;15:399–400.

