## Supplementary Note 1; Figure S1 for "scValue: value-based subsampling of large-scale single-cell transcriptomic data for machine and deep learning tasks"

### Supplementary Note 1: Experimental setup of the cell-type annotation tasks

scValue was benchmarked against six existing subsampling (sketching) methods, i.e., Uniform, GeoSketch, Sphetcher, Hopper, KH, and scSampler, across four cell-type annotation tasks that paired previously studied single-cell RNA sequencing (scRNA-seq) datasets with various machine or deep learning models. Below, we detail the experimental setup, including data information, reference-query splitting, sketching procedures, and training parameters.

**PBMC with scANVI**

We used the Peripheral Blood Mononuclear Cells (PBMC) dataset [1], which comprises 31,021 cells across 10 cell types with 2,000 highly variable genes (HVGs). This dataset was used to training the variational autoencoder-based scANVI model [2] in the previous study [3]. Following this study, we split PBMC into:

- a reference (training) set of 20,303 cells from SeqWell, DropSeq, and Indrop sequencing protocols.
- a query (test) set of 16,941 cells from the 10X V2 protocol.

Each of the seven sketching methods was applied to generate sketches of the reference set at varying percentages of 2%, 4%, 6%, 8%, and 10%, respectively, while the query set remained unchanged. Each resulting sketch was used to train a scANVI model following the official tutorial (https://docs.scarches.org/en/latest/scanvi_surgery

_pipeline.html). To mitigate overfitting, we set vae.train() to 100 epochs, scanvae.train() to 10 epochs, and model.train() to 10. All other parameters were kept at their default values. The trained model was then used to infer cell types for the full query set. A separate model was trained on the entire reference set to quantify performance loss due to sketching. Each training-and-annotation run was repeated 10 times (random seeds 42 to 51), and the annotation accuracy was recorded for each run.

**mBrain with scPoli**

The second task employed the mBrain dataset [4], which contains 56,399 cells across 10 cell types and 2,000 HVGs. This dataset includes cells from four independent studies and has been investigated in conjection with the variational autoencoder-based scPoli model [5]. We split mBrain into:

- a reference set: Saunders et al. (34,502 cells) and Zeisel et al. (7,394 cells)
- a query set: Schaum et al. (7,856 cells) and Rosenberg et al. (6,647 cells)

All seven sketching methods were applied to the reference set at 2%, 4%, 6%, 8%, and 10% of its size, with the query set left unchanged. For each sketch, we trained scPoli following the official tutorial (https://docs.scarches.org/en/latest/scpoli_surgery

_pipeline.html). When training the reference scPoli model, to prevent overfitting on smaller sketches, we set n_epochs to 5 and pretraining_epochs to 4 for 2-6% sketches; then we set n_epochs to 10 and pretraining_epochs to 8 for 8% and 10% sketches. When training the reference mapping scPoli model for the full query set, we set n_epochs to 10 and pretraining_epochs to 8. The learning rate was set to 0.01, leaving other parameters at defaults. A model trained on the full reference set (with n_epochs=10 and pretraining_epochs=8 for both the reference and mapping models) served as a baseline for evaluating the performance drop due to sketching. All experiments were repeated 10 times (random seeds 42 to 51), with annotation accuracies recorded for each run.

**CxGmin with CellTypist**

The third task concerned the logistic regression-based CellTypist model [6] and the CxG_min dataset [7]. Downloaded from the GitHub page of the study “scTab: Scaling cross-tissue single-cell annotation models” [7], this dataset is a minimal subset of the 22.2 million human CELLxGENE census data. It already includes training and test partitions, drawn from 70% and 15% of the human donors, respectively. Each partition comprises 32,768 cells across 164 cell types with 19331 protein-coding genes. We selected 3,000 HVGs to compute the top 50 principal components as input features for the sketching methods.

Again, we produced sketches of the reference set at 2%, 4%, 6%, 8%, and 10%, keeping the test set unchanged. Each sketch was used to train a CellTypist model according to the official tutorial (https://colab.research.google.com/github/Teichlab/cel

ltypist/blob/main/docs/notebook/celltypist_tutorial_cv.ipynb). Following the tutorial’s recommendation, we enabled feature selection during model training and left all parameters at defaults. A model trained on the full reference set served as the baseline for performance. Each experiment was repeated 10 times (random seeds 42 to 51), and the annotation accuracy was documented for each run.

**mACA with ACTINN**

The final task paired the neural networks-based ACTINN model [8] and the mACA dataset [9] consisting of 356k cells across 197 cell types with 20,116 genes. This model-dataset combination has been explored in the previous study [10]. Following this, we first filtered out cells with missing (“nan”) cell type labels and then split the remaining cells based on timepoints into

- a reference (training) set: 43,196 cells from mouse samples aged 18 months
- a query (test) set: 140,743 cells obtained from mouse samples aged 1, 3, 21, 24, or 30 months

The reference set was subsampled at 2%, 4%, 6%, 8%, and 10% with each of the seven sketching methods, while the query set remained intact. We trained ACTINN on each sketch using the default parameters (50 epochs, learning rate of 0.0001, minibatch size of 128). A model trained on the full reference set was also included for baseline comparison. All experiments were repeated 10 times (random seeds 42 to 51), and the resulting annotation accuracies were recorded.

### Supplementary figure

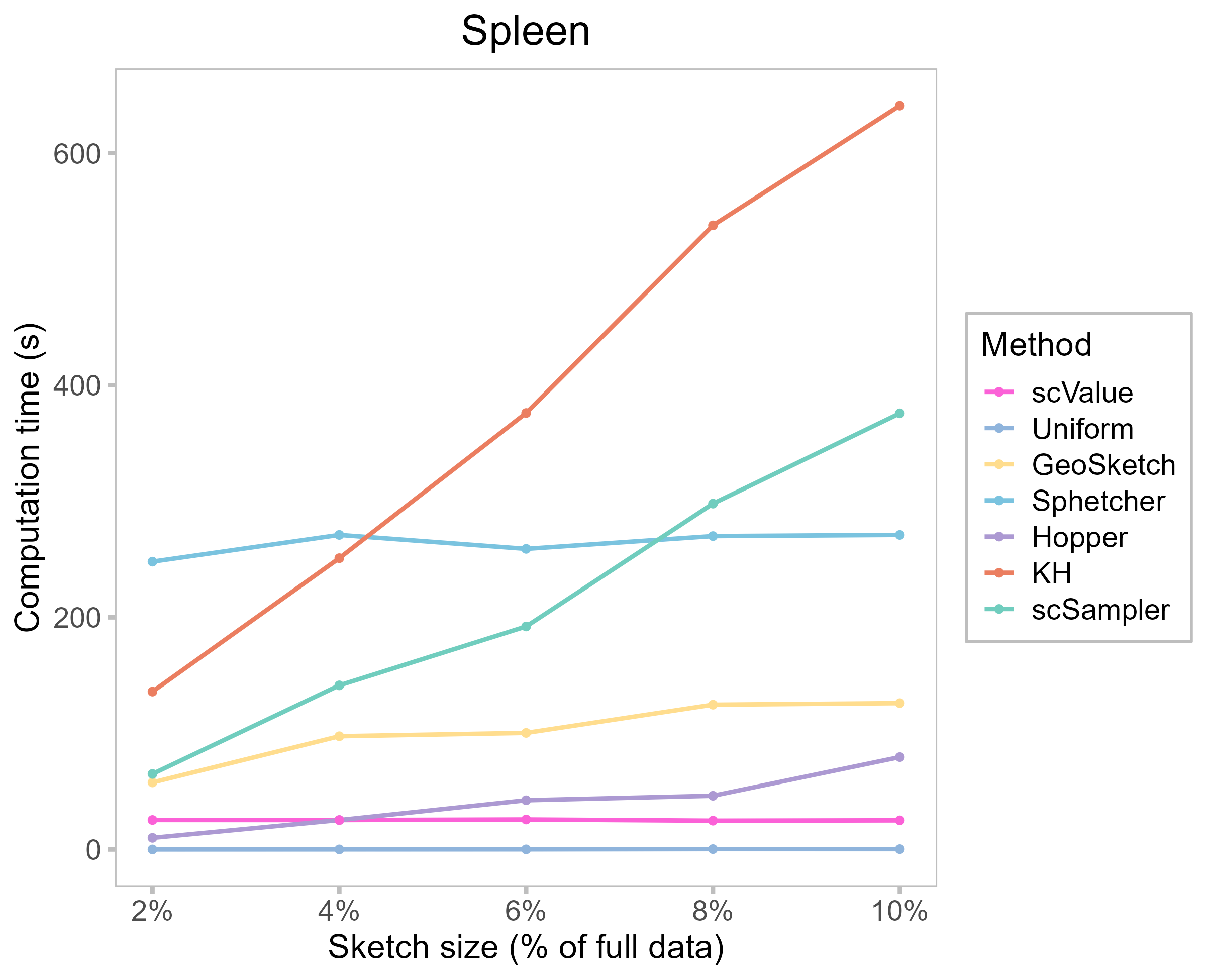

**Figure S1.** The Spleen dataset was used as an example to illustrate how computation time of each sketching methods changes as the sketch size increases from 2% to 10%. scValue’s computational time is nearly independent of the sketch size, which resonates with *O*(*BdN*log*N*) complexity.

### References

1.	Abdelaal T, Michielsen L, Cats D et al. A comparison of automatic cell identification methods for single-cell RNA sequencing data, Genome Biol 2019;20:194.

2.	Xu C, Lopez R, Mehlman E et al. Probabilistic harmonization and annotation of single-cell transcriptomics data with deep generative models, Mol Syst Biol 2021;17:e9620.

3.	Du ZH, Hu WL, Li JQ et al. scPML: pathway-based multi-view learning for cell type annotation from single-cell RNA-seq data, Commun Biol 2023;6:1268.

4.	Lotfollahi M, Naghipourfar M, Luecken MD et al. Mapping single-cell data to reference atlases by transfer learning, Nat Biotechnol 2022;40:121-130.

5.	De Donno C, Hediyeh-Zadeh S, Moinfar AA et al. Population-level integration of single-cell datasets enables multi-scale analysis across samples, Nature Methods 2023;20:1683-1692.

6.	Dominguez Conde C, Xu C, Jarvis LB et al. Cross-tissue immune cell analysis reveals tissue-specific features in humans, Science 2022;376:eabl5197.

7.	Fischer F, Fischer DS, Mukhin R et al. scTab: scaling cross-tissue single-cell annotation models, Nature Communications 2024;15:6611.

8.	Ma F, Pellegrini M. ACTINN: automated identification of cell types in single cell RNA sequencing, Bioinformatics 2020;36:533-538.

9.	Tabula Muris C. A single-cell transcriptomic atlas characterizes ageing tissues in the mouse, Nature 2020;583:590-595.

10.	Chen J, Xu H, Tao W et al. Transformer for one stop interpretable cell type annotation, Nat Commun 2023;14:223.
